# Staufen blocks autophagy in neurodegeneration

**DOI:** 10.1101/659649

**Authors:** Sharan Paul, Warunee Dansithong, Mandi Gandelman, Karla P. Figueroa, Tao Zu, Laura P.W. Ranum, Daniel R. Scoles, Stefan M. Pulst

## Abstract

**Objective:** The mechanistic target of rapamycin (mTOR) kinase is one of the master coordinators of cellular stress responses, regulating metabolism, autophagy, and apoptosis. We recently reported that Staufen1 (STAU1), a stress granule (SG) protein, was overabundant in fibroblast cell lines from patients with spinocerebellar ataxia type 2 (SCA2), amyotrophic lateral sclerosis, frontotemporal degeneration, Huntington’s, Alzheimer’s, and Parkinson’s diseases as well as animal models, and patient tissues. STAU1 overabundance is associated with mTOR hyperactivation and links SG formation with autophagy. Our objective was to determine the mechanism of mTOR regulation by STAU1.

**Methods:** We determined STAU1 abundance with disease- and chemical-induced cellular stressors in patient cells and animal models. We also used RNA binding assays to contextualize STAU1 interaction with *MTOR* mRNA.

**Results:** STAU1 and mTOR were overabundant in BAC-*C9orf72, ATXN2*^*Q127*^, and *Thy1*-TDP-43 transgenic mouse models. Reducing STAU1 levels in these mice normalized mTOR levels and activity and autophagy-related marker proteins. We also saw increased STAU1 levels in HEK293 cells transfected to express C9orf72-relevant dipeptide repeats (DPRs). Conversely, DPR accumulations were not observed in cells treated by *STAU1* RNAi. Overexpression of STAU1 in HEK293 cells increased mTOR levels through direct *MTOR* mRNA interaction, activating downstream targets and impairing autophagic flux. Targeting mTOR by rapamycin or RNAi normalized STAU1 abundance in a SCA2 cellular model.

**Interpretation:** STAU1 interaction with mTOR drives its hyperactivation and inhibits autophagic flux in multiple models of neurodegeneration. Staufen, therefore, constitutes a novel target to modulate mTOR activity, autophagy, and for the treatment of neurodegenerative diseases.

## Introduction

Autophagy dysfunction affects clearance of toxic aggregate-prone proteins in multiple neurodegenerative diseases (NDDs), including amyotrophic lateral sclerosis (ALS), frontotemporal degeneration (FTD), autism spectrum disorders (ASD), SCA2, Huntington’s, Alzheimer’s, and Parkinson’s diseases (HD, AD, and PD)^1-2^. NDDs caused by dominant mutations in human genes have been difficult to approach therapeutically, and few studies have targeted disease genes as the first step in pathogenesis. Additionally, stimulating autophagy by decreasing mTOR, using the pharmacologic mTOR inhibitor, rapamycin, or genetic reductions of mTOR, is beneficial for HD, FTD with ALS, ASD, and AD disease models^3-5^.

Analogous to other polyglutamine (polyQ) expansion neurodegenerative diseases [SCA types 1, 3, 6, 7, and 17; HD, spinal bulbar muscular atrophy (SBMA)], DNA CAG-repeat expansions in the *ATXN2* gene cause SCA2 and lead to formation of ATXN2-Q^exp^ protein aggregates^6,7^. Although the disease was initially described as cerebellar degeneration, it is now known to involve multiple neuronal subtypes outside of the cerebellum and can present as ALS, FTD with ALS, and PD^8^. Supporting ATXN2 as a therapeutic target for SCA2 and ALS, reducing ATXN2 levels in SCA2 or ALS-TDP-43 mouse models, using antisense oligonucleotides (ASOs) or genetic interaction, delayed neurodegeneration^9,10^.

There is considerable interest to identify other potential therapeutic targets for NDDs by searching for proteins interacting with disease-linked mutant genes. Using immunoprecipitation and mass spectrometry, we identified Staufen1 (STAU1) as an ATXN2 interactor^11^. STAU1 is a double-stranded RNA binding protein (dsRBP) and a component of stress granules (SGs) that can influence RNA stability and translation^12,13^. It can bind to a large number of mRNAs via the 3’UTR and initiate STAU-mediated decay (SMD), a process similar to nonsense-mediated decay^12,14-16^.

We previously demonstrated that STAU1 was overabundant in fibroblast (FB) cell lines from individuals with SCA2, HD, ALS-TDP-43, ALS/FTD-C9orf72, and AD as well as animal models of multiple NDDs. STAU1 overabundance was not transcriptional but tied to reduced autophagic degradation^11,17^. STAU1 is recruited to ATXN2-Q^exp^ aggregates in Purkinje cells (PC) in SCA2^11^. In cellular models, STAU1 overabundance amplified the proapoptotic activation of the unfolded protein response (UPR) and was sufficient to mediate caspase-3 cleavage^18^. Although reducing *Stau1* gene dosage *in vivo* decreased the burden of ATXN2-Q^exp^ aggregates and improved motor behavior of SCA2 mice with normalization of key PC proteins^11^, the role of STAU1 in autophagy and neurodegeneration remained poorly characterized.

Here we provide a mechanistic link for STAU1 in regulating mTOR. We determined that decreasing mTOR activity can normalize STAU1 abundance, and inversely, that decreasing STAU1 levels improves autophagic flux. Furthermore, we show that reducing STAU1 levels *in vivo* restores mTOR and autophagy pathway protein levels in brains of SCA2 and ALS-TDP-43 animal models and in a cellular model of C9orf72-DPR toxicity. We describe STAU1 overabundance as a common feature across multiple neurodegenerative diseases of varied origin suggesting a functional link among STAU1, mTOR signaling, inhibition of autophagic flux and neurodegeneration that could be exploited in therapeutic development.

## Materials and Methods

### DNA constructs

The plasmid constructs used in this study were 3XFlag-tagged STAU1 or His-tagged STAU1 or GFP^11^. pEGFP-LC3 (human) was a gift from Toren Finkel (Addgene, Plasmid #24920). pMXs GFP-LC3-RFP was a gift from Noboru Mizushima (Addgene, plasmid #117413). pAG416-Gal-PA50 (Addgene #84902) and pAG416-Gal-GA50 (Addgene #84903) were a gift from Aaron Gitler. RNA-binding domain 3 deletion of STAU1 (STAU1-RBDΔ3) constructs were generated using *STAU1* cDNAs as templates, respectively^11^. The PCR products were cloned into pCMV-3XFlag plasmid (Agilent Technologies, USA). To generate a GFP-LC3-RFP autophagic flux reporter construct, the GFP-LC3-RFP sequence from the pMXs-GFP-LC3-RFP plasmid was subcloned into pcDNA3-neomycin plasmid. For mammalian expression of DPR plasmids (poly-GA^50^ and poly-PA^50^), DPR sequences with Myc/Flag in-frame sequence were subcloned into pCMV-3XFlag plasmid at *BamH*I and *Xho*I site. For the 5’UTR-*MTOR*-Luciferase (*MTOR*^*5’UTR-WT*^) reporter assay, we subcloned the luciferase reporter gene from pGL4.10[Luc2] vector (Promega) into pcDNA3.1 (Thermo Fisher) at *Hind* III and *Xba* I sites, designated as CMV-LUC. 120 bp oligonucleotides upstream of the *MTOR* start codon were synthesized and ligated to upstream of the luciferase of CMV-LUC plasmid at *Hind* III site. 5’UTR of *PCP2* (262 bp) was PCR amplified from HEK293 cell cDNA library and cloned into CMV-LUC plasmid at *Hind* III site. To generate a mutant *MTOR*-5’UTRΔ-luciferase-(*MTOR*^*5’UTRΔ-Mt*^) construct, GCGGT and GGGGCCT sequences (33 and 45^th^ positions) from the predicted minimum free energy (MFE) secondary structure of 5’UTR of *MTOR* sequence [RNAfold WevServer: http://rna.tbi.univie.ac.at/cgi-bin/RNAWebSuite/RNAfold.cgi] were replaced with AAAAA and AAAAAAA in *MTOR* upstream oligonucleotides. All constructs were verified by sequencing. 3XFlag is referred to as Flag in the text and figures.

### siRNAs and reagents

The siRNAs used in this study were: All Star Negative Control siRNA (Qiagen, Cat# 1027280), human *siSTAU1*: 5’-CCUAUAACUACAACAUGAGdTdT-3’^15,16^ and *siMTOR:* 5’-GAG CCUUGUUGAUCCUUAA-3’^19^. All siRNA oligonucleotides were synthesized by Invitrogen, USA. The oligonucleotides were deprotected and the complementary strands were annealed. Bafilomycin A1 (InvivoGen USA, Cat# tlrl-baf1), Rapamycin (InvivoGen USA, Catalog # tlrl-rap), Thapsigargin (Tocris USA, Cat# 1138), Tunicamycin (Tocris USA, Cat# 3516), Ionomycin (Tocris USA, Cat# 1704) and Sodium arsenite solution (Sigma-Aldrich, Cat# 1062771000) were used in this study.

### Cell line authentication

In order to adhere with the NIH guideline on scientific rigor in conducting biomedical research on the use of biological and/or chemical resources (NOT-OD-15-103) we authenticated our cell lines utilizing STR analysis on 24 loci (GenePrint 24 system, https://www.promega.com/products/cell-authentication-sample-identification/mixed-sample-analysis/geneprint-24-system/?catNum=B1870).

### Cell culture and transfections

The following primary human fibroblasts were obtained from the Coriell Cell Repositories (Camden, NJ, USA): Normal FBs (#ND29510, #ND34769, and #ND38530), two FBs from patients at risk of frontotemporal degeneration (FTD) with *C9orf72* expansions (#ND42504 and #ND42506), All FBs, including ATXN2-Q22/58 (ATXN2-Q58) knock-in (KI) HEK293 cells^11^ used in this study, were maintained in DMEM medium containing 10% fetal bovine serum. All procedures for human FBs were approved by the Institutional Review Board (IRB) at the University of Utah.

For over-expression of recombinant proteins, HEK293 cells were plated on 6-well dishes and incubated overnight. The cells were then transfected with plasmid DNAs and harvested 48 hrs post-transfection and processed as two aliquots for protein and RNA analyses. For siRNA experiments, cells were transfected with siRNAs using lipofectamine 2000 transfection reagent (ThermoFisher Scientific) according to the manufacturer’s protocol. Prior standardization experiments showed that maximum silencing was achieved 4-5 days post-transfection. For luciferase reporter assays, HEK293 cells were cultured on 12-well plates and incubated overnight. Cells were then transfected using lipofectamine 2000 transfection reagent with the following plasmid constructs: Flag-tagged STAU1 or STAU1^RBDΔ3^ or Flag plasmids with the wildtype or mutant *5’UTR-MTOR-LUC* or luciferase empty vector constructs and Renilla luciferase plasmid (pRL-SV40 vector, Promega) according to an experimental set-up. 48 h post-transfection, the cells were subjected to the luciferase assay using the Duel-Glo Luciferase Assay System following the manufacturer’s protocol (Promega). For autophagy activity assays, HEK293 and ATXN2-58 KI cells were stably transfected with GFP-LC3 or GFP-LC3-RFP plasmids and selected and maintained with G418 (Geneticin) (Invitrogen) or puromycin. GFP-LC3 stably expressing cells were seeded on 6-well dishes and incubated overnight. The cells were then treated with bafilomycin A1 (Baf) for 6 hr and harvested followed by western blot analyses. For GFP-LC3-RFP reporter assays, cells stably expressing GFP-LC3-RFP were plated on 96-well plates (20,000 cells/well). Following overnight incubation, the cells were treated with rapamycin for 24 hr, and fluorescence was measured with the Beckman Coulter DTX 880 Multimode Detector and settings: GFP (excitation 485 nm and emission 535 nm) and RFP (excitation at 550 nm and emission at 595 nm).

### Mice

*ATXN2*^*Q127*^ (*Pcp2*-ATXN2[Q127]) mice^20^ were maintained in a B6D2F1/J background. The *Stau1*^*tm1Apa(-/-)*^ (*Stau1*^*-/-*^) mouse^21^ was a generous gift from Prof. Michael A. Kiebler, Ludwig Maximilian University of Munich, Germany, and maintained in a C57BL/6J background. *ATXN2*^*Q127*^ mice were crossed with *Stau1*^*-/-*^ mice to generate *ATXN2*^*Q127*^/*Stau1*^*+/-*^ and *ATXN2*^*WT*^/*Stau1*^*+/-*^ mice. These mice were then interbred to generate *ATXN2*^*Q127*^/*Stau1*^*-/-*^ and *ATXN2*^*WT*^/*Stau1*^*-/-*^ mice. These mice were in a mixed background of B6D2F1/J C57BL/6J. B6;SJL-Tg(Thy1-TARDBP)4Singh/J (Stock 012836)^22^ purchased from Jackson Laboratories. Jackson Laboratories B6SJLF1/J mice (Stock No. 100012) were backcrossed to C57BL/6J to N2 before crossing them to *Stau1*^*-/-*^ mice. Genotyping of animals was accomplished according to published protocols^20-22^. All mice were bred and maintained under standard conditions consistent with National Institutes of Health guidelines and conformed to an approved University of Utah IACUC protocol. Wildtype and BAC-*C9orf72* (C9-500) mouse^23^ brain extracts were prepared by the LPWR laboratory and shipped to the SMP laboratory.

### Primary culture of cortical neurons

Primary cortical neuron cultures were prepared from neonatal wild-type or *STAU1*^*-/-*^ mice [*Stau1*^*tm1Apa(-/-)*^]^21^. Brain cortices from 6-7 animals were isolated and incubated with 50 units of papain (Worthington Biochemical, USA) in Earle’s balanced salt solution (EBSS) with 1.0 mM L-cysteine and 0.5 mM EDTA for 15 min. at 37°C, followed by washing in EBSS and mechanical trituration. Remaining tissues were removed by filtration through a 50 μm strainer (Falcon). Neurons were seeded at 50×10^3^ per cm^2^ on plates coated with poly-L-ornithine and laminin in Neurobasal Plus medium containing 2% B27 Plus supplement (Life Technologies, USA). On day 2, 10 μM cytosine arabinoside was added for 24 hr to prevent proliferation of glial cells, and 75% of culture medium volume was replenished every 2-3 days from thereon. Experiments were conducted on days 9-10 by replacing all culture mediums with fresh one containing thapsigargin or vehicle (DMSO), and after 18 hr protein lysates were prepared following standard procedures.

### Antibodies

The antibodies used for western blotting and their dilutions were as follows: rabbit anti-Staufen antibody [(1:5000), Novus biologicals, NBP1-33202], LC3B Antibody [(1:7000), Novus biologicals, NB100-2220], monoclonal anti-FLAG M2 antibody [(1:10,000), Sigma-Aldrich, F3165], monoclonal anti-β-Actin−peroxidase antibody (clone AC-15) [(1:30,000), Sigma-Aldrich, A3854], SQSTM1/p62 antibody [(1:4000), Cell Signaling, Cat# 5114], mTOR antibody [(1:4000), Cell Signaling, Cat# 2972], Phospho-mTOR (Ser2448) antibody [(1:3000), Cell Signaling, Cat# 2971], Phospho-p70 S6 Kinase (Thr389) antibody [(1:3000), Cell Signaling, Cat# 9205], 6x-His Tag Monoclonal Antibody (HIS.H8), HRP [(1:10,000) (ThermoFisher Scientific, MA1-21315-HRP)], Sheep-anti-Digoxigenin-POD, Fab fragments [(1:10,000), Roche Life Science, Cat# 11207733910], GAPDH (14C10) rabbit mAb [(1:30,000), Cell Signaling, Cat# 2118], The secondary antibodies were: Peroxidase-conjugated AffiniPure goat anti-rabbit IgG (H + L) antibody [(1:5000), Jackson ImmunoResearch Laboratories, Cat# 111-035-144] and Peroxidase-conjugated horse anti-mouse IgG (H+L) antibody [(1:5,000) (Vector laboratories, PI-2000)].

### Preparation of protein lysates and western blotting

Cellular extracts were prepared by a single-step lysis method^11,17^. The harvested cells were suspended in SDS-PAGE sample buffer [Laemmli sample buffer (Bio-Rad, Cat# 161-0737)] and then boiled for 5 min. Equal amounts of the extracts were used for western blot analyses. Mice were deeply anesthetized with isoflurane. Mouse cerebella or spinal cord were removed and immediately submerged in liquid nitrogen. Tissues were kept at −80°C until the time of processing. Mouse cerebellar or spinal cord protein extracts were prepared by homogenization of tissues in extraction buffer [25 mM Tris-HCl pH 7.6, 300 mM NaCl, 0.5% Nonidet P-40, 2 mM EDTA, 2 mM MgCl_2_, 0.5 M urea and protease inhibitors (Sigma-Aldrich, P-8340)] followed by centrifugation at 4°C for 20 min at 14,000 RPM. Only supernatants were used for western blotting. Protein extracts were resolved by SDS-PAGE and transferred to Hybond P membranes (Amersham Bioscience, USA), and then processed for western blotting according to our published protocol^11,17^. Immobilon Western Chemiluminescent HRP Substrate (EMD Millipore, Cat# WBKLSO500) was used to visualize the signals, which were detected on the ChemiDoc MP imager (Bio-Rad). For some blots, we used a film developing system and band intensities were quantified by ImageJ software analyses after inversion of the images. Relative protein abundances were expressed as ratios to ACTB or GAPDH.

### Immunoprecipitations

To determine protein-RNA interactions, protein-RNA immunoprecipitation (IP) experiments were carried out using endogenous STAU1 IP. The preparation of whole-cell extracts and IP methods followed previously published methods^11,24^. Briefly, whole-cell extracts were prepared using the two-step lysis procedure with cytoplasmic extraction buffer [25 mM Tris-HCl pH 7.6, 10 mM NaCl, 0.5% NP40, 2 mM EDTA, 2 mM MgCl_2_, and protease and RNase inhibitors], and nuclear lysis buffer [25 mM Tris-HCl, pH 7.6, 500 mM NaCl, 0.5% Nonidet P-40, 2 mM EDTA, 2 mM MgCl_2_, and protease and RNase inhibitors]. The nuclear extracts were combined with the cytoplasmic extracts and denoted as whole-cell extracts. Specifically, while combining cytoplasmic and nuclear extracts, the NaCl concentration was adjusted to physiologic buffer conditions (∽150 mM) to preserve *in vivo* interactions. For endogenous STAU1 IP, equal amount of rabbit anti-Staufen antibody (Novus Biologicals, NBP1-33202) and normal rabbit IgG (Cell Signaling, Cat# 2729) (5.0 μg each), as control, immobilized with Pierce™ Protein A/G agarose beads (ThermoFisher Scientific, Cat# 20421) according to the manufacturere’s protocol. Equal amounts of whole-cell extracts were incubated with STAU1 or control antibody-coupling agarose beads for overnight at 4°C. The beads were then washed several times with a washing buffer (10 mM Tris-HCl, pH 7.6, 10 mM HEPES, 200 mM NaCl, 2.0 mM MgCl_2_, 0.5 mM EDTA, 0.05% Nonidet P-40, 10% glycerol, and protease and RNase inhibitors). The halves of STAU1 or control beads were analyzed by western blotting to validate IP. Total RNAs were extracted from the other halves of the beads using TRIzol™ Reagent (ThermoFisher Scientific, Cat# 15596026) according to manufacturer’s protocol. RNAs were used to synthesize cDNAs using the ProtoScript cDNA synthesis kit (New England Biolabs, USA) and subjected to RT-PCR analyses to determine STAU1-RNA interactions. Primer pairs designed for RT-PCR are given as forward and reverse, respectively, GAPDH-F: 5’-ACATCGCTCAGACACCATG-3’, GAPDH-R: 5’-TGTAGTTGAGGTCAATGAAGGG-3’, MTOR-F: 5’-CGAACCTCAGGG CAAGATG-3’ and MTOR-R: 5’-TTTCCTCATTCCGGCTCTTTAG-3’.

### In vitro RNA binding (Northwestern) assay

Northwestern blot assays were performed to determine STAU1 and *MTOR-5’UTR* interaction following a published protocol with some modifications^11^. BL21<DE3> cells carrying His-STAU1, His-GFP, and empty pET vectors, were grown to mid-log phase. Whole-cell lysates from harvested cells were run on SDS-PAGE and electro-blotted onto a Hybond P membrane. The transferred proteins were re-natured as follows: the blot was first incubated with binding buffer (0.1 M HEPES, pH 7.9, 0.1 M MgCl_2_, 0.1 M KCl, 0.5 μM ZnSO_4_) with 1 mM DTT and 6 M urea for 5 min at room temperature. The blots were then incubated, for 5 min each, through five serial twofold dilutions of urea with binding buffer/1 mM DTT with continuing incubation steps until the binding buffer was 1 mM DTT without urea. Blot pre-hybridizations were carried out with binding buffer (1 mM DTT, 5% BSA, 1 μg/ml yeast tRNA) for 1 hr at room temperature. The blots were then hybridized with DIG-labelled RNA probes in binding buffer (1 mM DTT, 0.25% BSA, 1 μg/ml yeast tRNA) overnight at 4°C. After several washes with 0.1% Tween 20/PBS, the membranes were incubated with anti-DIG-POD antibody in 4% skim milk in 0.1% Tween 20/PBS for 2 hr at room temperature. Following three additional washes with 0.1% Tween 20/PBS, signals were detected by using the Immobilon Western Chemiluminescent HRP Substrate according to the manufacturer’s protocol.

### Statistical analysis

Two-tailed *t*-tests or one-way ANOVA followed by Bonferroni’s multiple comparisons test were used to determine significant differences among groups. The levels of significance were indicated as follows: \**P* ≤ 0.05, \*\**P* ≤ 0.01, \*\*\**P* ≤ 0.001 and ns= *P* > 0.05. Means ± SDs are presented throughout unless otherwise specified. Statistical tests were carried out using GraphPad Prism version 8.

## Results

### STAU1 levels are increased in neurodegenerative diseases

We previously demonstrated that STAU1 overabundance in neurodegenerative disease FBs, mouse models, and human tissues associate with autophagy malfunction by way of mTOR hyperactivation^17^. To evaluate the role of STAU1 in neurodegeneration with emphasis on mTOR and autophagy, we used a well-characterized ALS/FTD BAC-C9orf72 mouse line expressing a human BAC with an expanded G_4_C_2_ tract 500 repeats in length (BAC-*C9orf72*)^23^. By western blot analysis, BAC-*C9orf72* mouse brain extracts demonstrated greatly increased Stau1 levels similar to those seen in *ATXN2*^*Q127*^, and ALS-TDP-43 mice (**Fig 1A, B**)^17^. Associated with elevated Stau1, we found that autophagy pathway components were altered predicting reduced flux by inefficient autophagosome-lysosome fusion. This included total and phosphorylated mTor, phospho-p70 S6 kinase (P-S6K) as well as p62/SQSTM1 and LC3-II, which functions as a selective autophagy receptor for degradation of ubiquitinated substrates.

**Figure 1.**
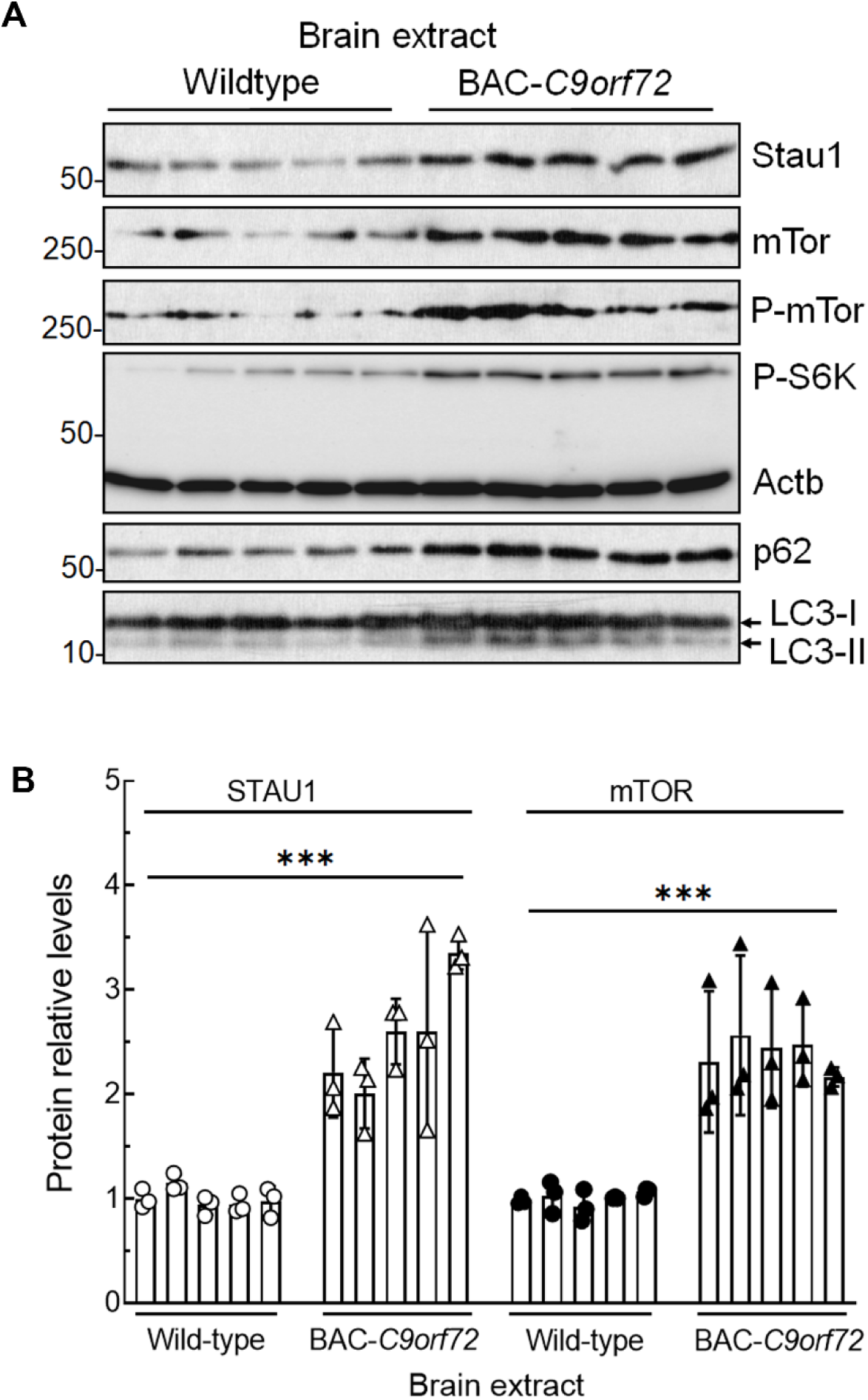
Central nervous system (CNS) tissues from mice with *C9orf72* mutations or expansions have increased Stau1 levels, abnormal mTor activity and autophagy defects. (**A, B)** Western blot analyses showed increased Stau1, mTor, P-mTor, P-S6K, p62 and LC3-II levels in and BAC-*C9orf72* mouse brain extracts (20 wks of age; 5 animals per group) compared to wildtype controls. Stau1 and mTor protein levels were normalized to Actb and quantified average fold changes for Stau1 and mTor are shown below the blots in (**B)**. Each lane represents an individual mouse. Actb was used as a loading control and the blots are from three replicate blots. One-way ANOVA followed by Bonferroni’s multiple comparisons test. Data are mean ± SD, ns = *P* > 0.05, *\*\*\*P* < 0.001

### STAU1 responds to a variety of stressors

Accumulating evidence indicates that stress-induced intracellular changes of endoplasmic reticulum (ER) and unfolded protein response (UPR) functions lead to neurotoxicity in a number of NDDs^25^. We wanted to test whether subjecting cells to various forms of stress would affect STAU1 and autophagic functions similarly to mutation or overexpression of disease-linked genes^17^. We used HEK293 cells and subjected them to endoplasmic reticulum (ER) stressors (thapsigargin or tunicamycin), a calcium ionophore (ionomycin), oxidative stress (tunicamycin), metal stressor (sodium arsenite), and temperature stress (hyperthermia) followed by western blotting. The different forms of stress all induced STAU1 overabundance, mTOR elevation, and increased phosphorylation of mTOR targets (**Fig 2A-E**). To confirm the involvement of STAU1 in the activation of this pathway, we established primary cultures of cortical neurons from wildtype and *Stau1*^-/-^ mice^21^. For wildtype neurons, treatments with thapsigargin induced Stau1 overabundance, activated mTor, and increased p62 and LC3-II levels. In contrast, *Stau1*^-/-^ neurons showed minimal response to thapsigargin (**Fig 2F**), indicating that the activation of mTOR and downstream targets were STAU1-dependent. Although the precise signaling pathways are not yet known, these findings indicate that STAU1 is positioned at the crossroads of several cellular stress pathways.

**Figure 2.**
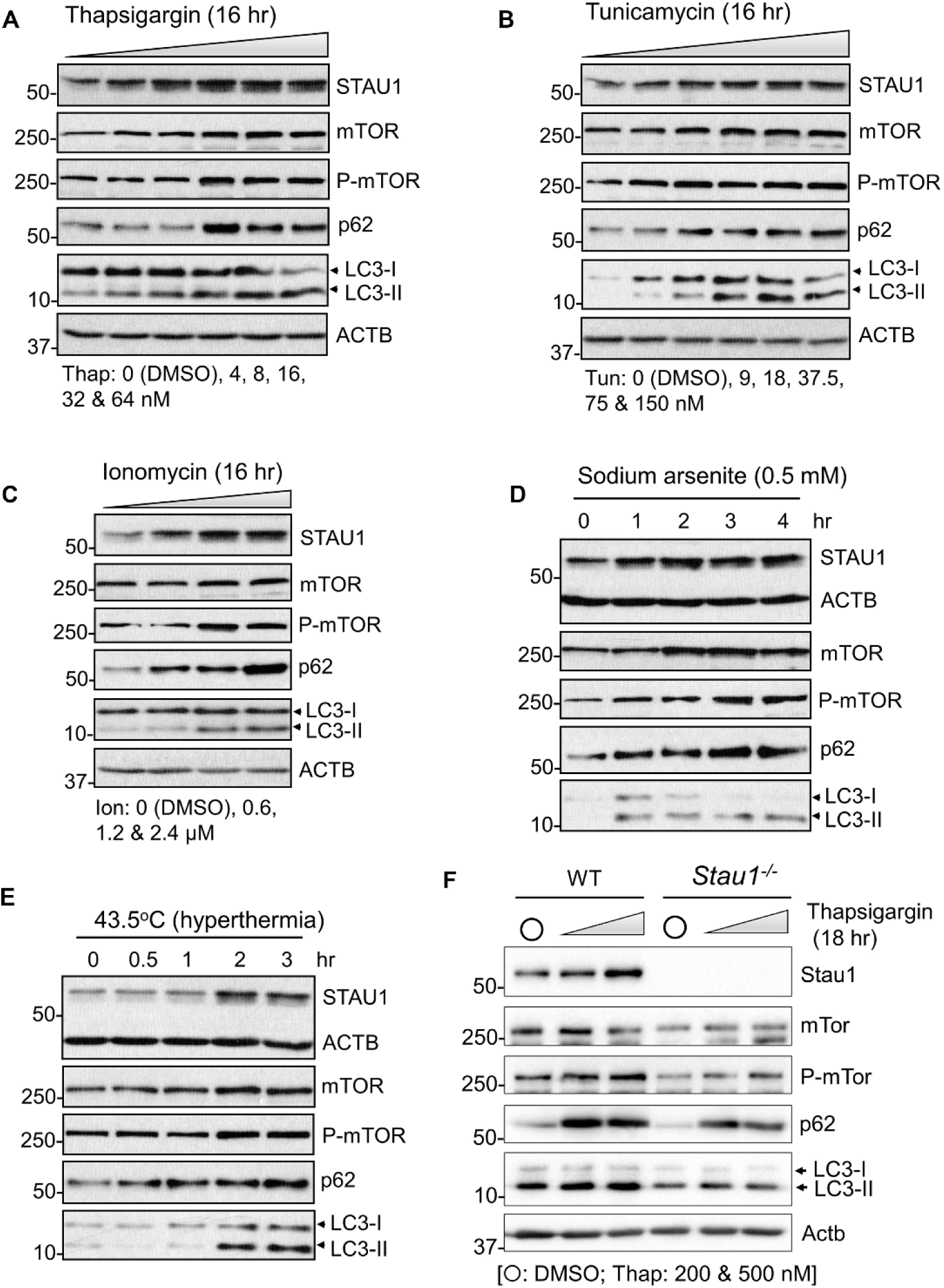
STAU1 responds to a variety of stressors. **(A-E)** HEK293 cells were treated with the indicated stressors and analyzed by western blotting. The treatments increased STAU1, mTOR and P-mTOR levels, and reduced autophagy activity (increased p62 and LC3-II levels). **(F)** Mouse cortical neurons treated with thapsigargin also showed increased Stau1, mTor, P-mTor, p62, and LC3-II (left), but neurons null for Stau1 are resistant to thapsigargin-mediated induction of these genes (right). ACTB/Actb was used as loading control. Representative blots of three independent experiments are shown.

### STAU1 binds mTOR mRNA enhancing translation

We previously showed that exogenous expression of STAU1 in HEK293 cells resulted in the same alterations that were seen in cells with NDD-linked mutations^17^. Expression of STAU1 in wildtype cells resulted in increased protein levels of mTOR, P-mTOR, p62, and LC3-II **(Fig 3A)**. These results indicated that STAU1 overabundance was sufficient to drive autophagic dysfunction. Notably, mTOR transcripts were found to be unchanged upon STAU1 overexpression in HEK293 cells suggesting that mTOR protein changes were potentially related to changes in translation^17^.

**Figure 3.**
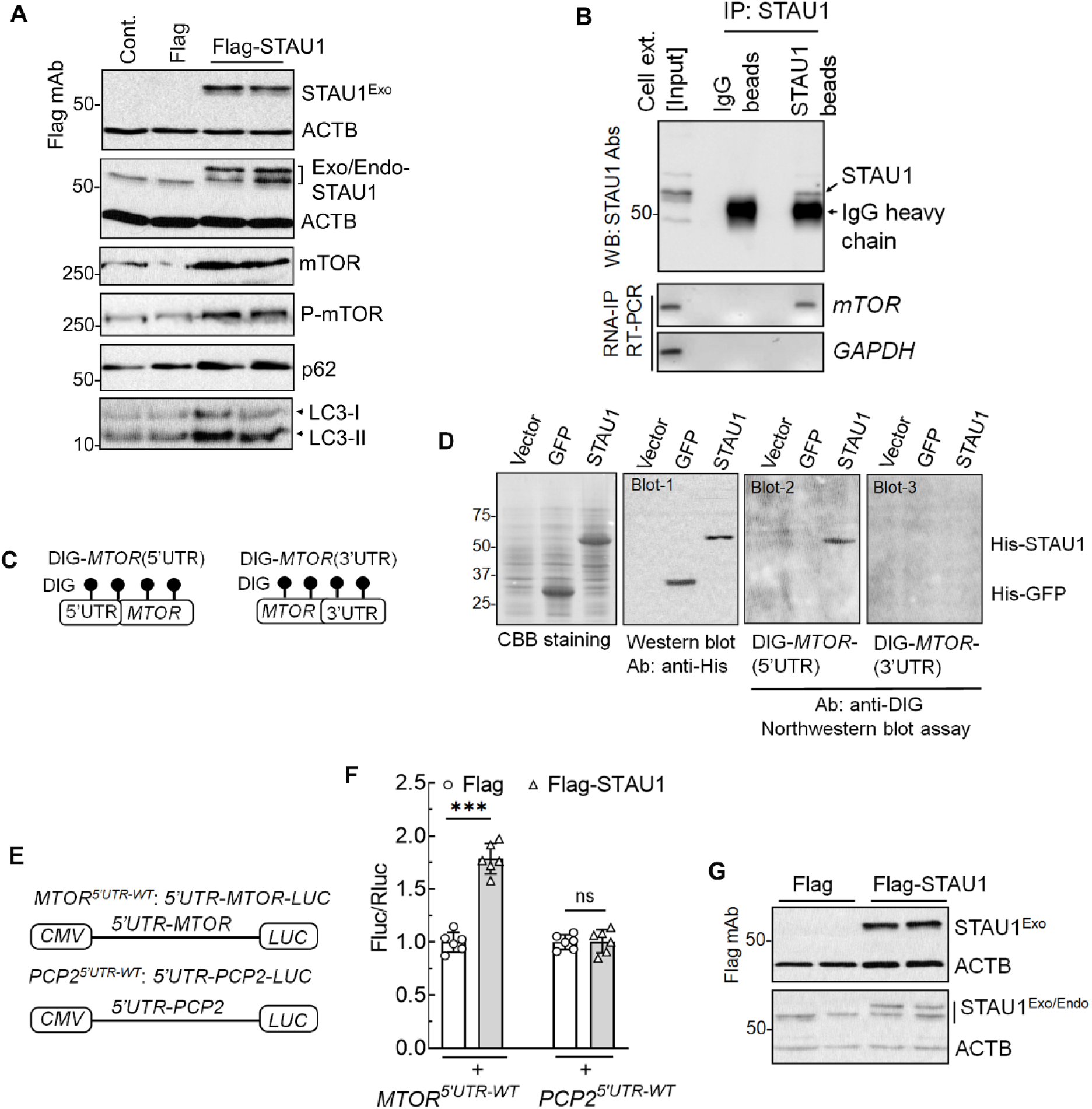
STAU1 interaction with the 5’UTR of *MTOR* enhances *MTOR* translation and mTOR signaling. **(A)** STAU1 overexpression elevates mTOR. HEK293 cells exogenously expressing Flag-tagged STAU1 were analyzed 48 hrs post-transfection by western blotting and showed increased levels of mTOR, P-mTOR, p62 and LC3-II. (**B-G)** STAU1 regulates mTOR expression through binding to *MTOR-5’UTR*. **(B)** Non-RNase A treated HEK293 whole-cell extracts were subjected to protein-RNA IP using anti-STAU1 antibody-coupling agarose beads. After washing with wash buffer (200 mM NaCl), the protein-RNA bound beads were analyzed by western blotting and RT-PCR showing STAU1 pulled down *MTOR* RNA, but not control *GAPDH*. **(C, D)** STAU1 binds directly to human *MTOR* RNA in a manner requiring the 5’UTR by Northwestern blot analysis. (**C)** Schematic of DIG-labeled human *MTOR* RNA probes with 5’UTR and 3’UTR sequences. **(D)** STAU1 directly interacts with *5’UTR-MTOR* RNA (blot 2), but not with *MTOR-3’UTR* RNA (blot 3). Bacterially expressed His-STAU1, His-GFP, or control bacterial lysate were stained with Coomassie brilliant blue (CBB) on SDS-PAGE or western blotted with anti-His antibody (blot 1). Protein blots were hybridized with DIG-labelled *MTOR* RNA probes followed by anti-DIG antibody staining. His-GFP shows no interactions. **(E-G)** Schematic of *5’UTR-MTOR-LUC* and *5’UTR-PCP2-LUC* (control) reporters (**E**). Exogenous STAU1 expression induces increased expression of the *5’UTR-MTOR-LUC* construct but not *5’UTR-PCP2-LUC* (*PCP2-3’UTR* substrate for SMD^11^ on luciferase reporter assay (**F**). Western blotting showing expression of STAU1 (**G**). ACTB was used as a loading control; blots are from three replicate experiments. Ordinary one-way ANOVA followed by Bonferroni’s multiple comparisons test. Data are mean ± SD, ns = *P* > 0.05, \*\*\**P* < 0.001

As STAU1 can enhance translation of mRNAs upon binding to the 5’UTR of the respective transcripts^26^, we tested whether STAU1 was able to bind *MTOR* mRNA and whether this interaction increased its translation. We employed two independent approaches to establish an interaction between STAU1 and *MTOR* mRNA. First, we used STAU1 protein-RNA immunoprecipitation (IP) in HEK293 cell extracts followed by RT-PCR and demonstrated that endogenous STAU1 pulled down *MTOR RNA* but not control *GAPDH* mRNA (**Fig 3B**). Second, northwestern blotting experiments were performed using bacterially expressed recombinant His-tagged STAU1 protein and *in vitro* transcribed DIG-labelled *MTOR* RNA probes. We generated probes: DIG-*MTOR*(5’UTR) and DIG-*MTOR*(3’UTR) (**Fig 3C**), which allowed us to identify those parts of the *MTOR* RNAs that interacted with STAU1. STAU1 showed binding to *MTOR*(5’UTR) (blot 2) RNA, but not with *MTOR*(3’UTR) RNA (blot 3), thereby showing the specificity of the interaction (**Fig 3D**). Thus, STAU1 interactions with 5’UTRs of *MTOR* mRNAs suggest its role for translation.

To test the consequence of STAU1 binding to *MTOR* mRNA, we performed luciferase reporter assays with constructs of *MTOR*-5’UTR-*LUC* and *PCP2*-5’UTR-LUC under control of the *CMV* promoter. *PCP2* mRNA is known to interact with STAU1 via its 3’UTR triggering SMD^17^. Exogenous expression of STAU1 in HEK293 cells resulted in significant induction of luciferase activity only from the *MTOR*-5’UTR-*LUC* construct (**Fig 3E-G**).

We next analyzed STAU1 function in the regulation of mTOR expression in a cell culture model with emphasis on *STAU1* RNA binding domain and *MTOR* RNA structures. Together with other published observations and our data demonstrate that STAU1 can bind to 5’UTRs of specific mRNAs to regulate their expression (**Fig 3)**^26^, The dsRBD3 of STAU1 is essential and acts as the major functional domain for target RNA interaction and its expression^27-29^. We used STAU1 with dsRBD3 deletion mutant by which STAU1 regulates mTOR expression. HEK293 cells were transfected with STAU1 constructs alone or co-transfected with *5’UTR-MTOR-LUC* reporter construct followed by western blotting or luciferase assay. In both assays, full-length STAU1, but not STAU1 with a deleted dsRBD3, resulted in enhanced translation of the *MTOR m*RNA (**Fig 4A-D**). Thus, these data suggest that STAU1 with dsRBD3 is essential to regulate the *MTOR* expression via its 5’UTR, corroborating STAU1 mediated translational efficiency^26^. The STAU1 dsRBD3 domain mediates interactions with specific secondary structure RNA stem-loops^27-29^. We, therefore, introduced unpaired bases in the *MTOR*-5’UTR in order to disrupt predicted stem-loop structures. When co-expressed with STAU1, the *MTOR-*5’UTR*-LUC* construct with small deletions in HEK293 cells resulted in significantly reduced luciferase activity (**Fig 4E-H**).

**Figure 4.**
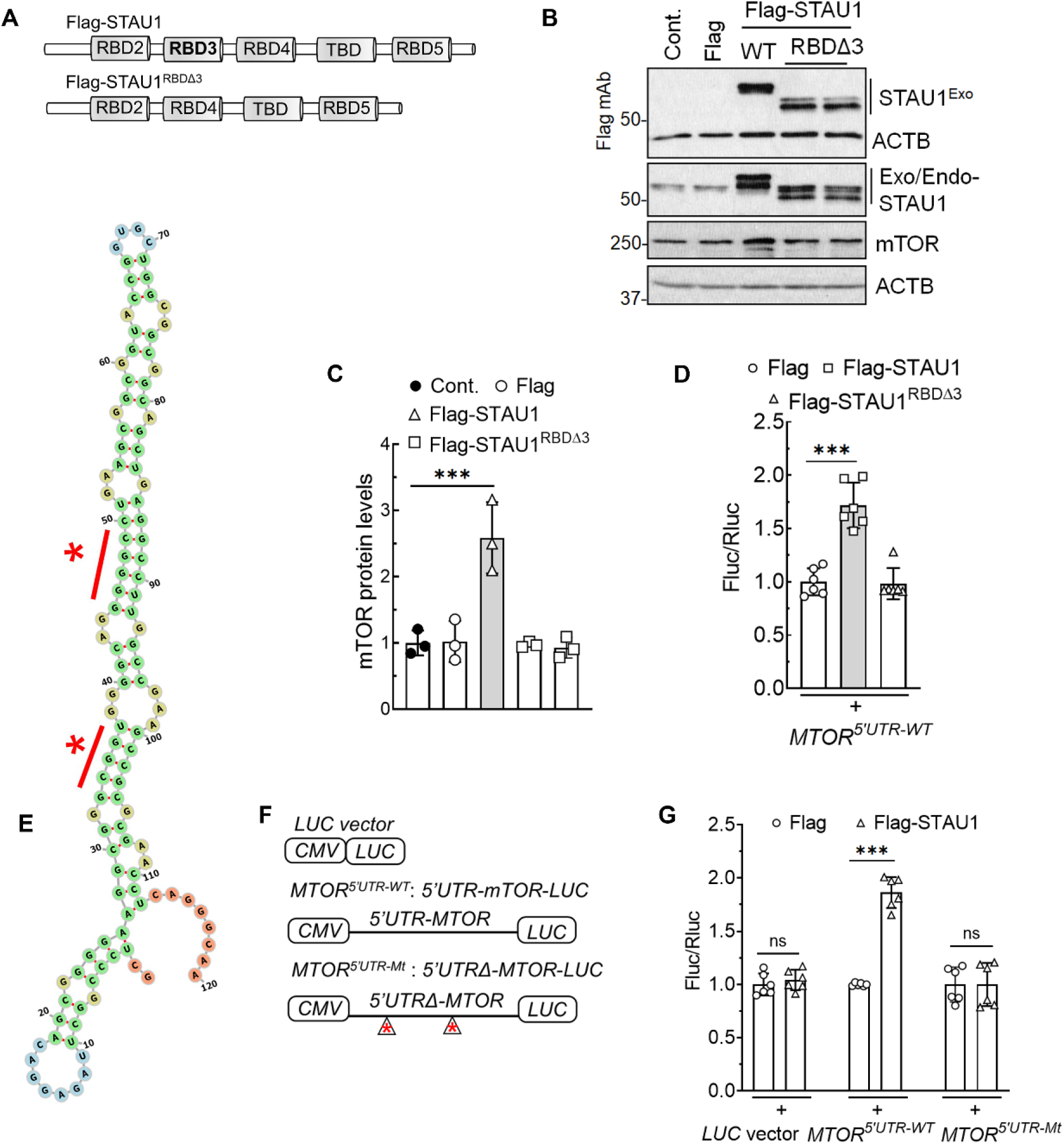
STAU1 induces *MTOR* mRNA translation. **(A-D)** The RNA binding domain 3 (RBD3) of STAU1 is required to induce mTOR expression. Schematic representation of STAU1 and mutant STAU1 excluding RBD3 (STAU1^RBDΔ3^) **(A)**. HEK293 cells were transfected with STAU1 constructs alone or co-transfected with *5’UTR-MTOR-LUC* reporter construct followed by western blotting or luciferase assay **(B-D)**. *S*ignificantly increased endogenous mTOR level (**B, C**) or luciferases activity (**D**) are achieved only through the STAU1 construct retaining the RBD3. (**E-G)** STAU1 increases translation of *MTOR* mRNA bearing double-stranded RNA stem-loop. *MTOR 5’UTR* secondary structure and asterisk represents changes of bases (**E**). (**F, G)** Depicted are empty Luciferase (LUC), wildtype *5’UTR-MTOR-LUC* (*MTOR*^*5’UTR-WT*^), or mutant *5’UTRΔ-MTOR-LUC* (*MTOR*^*5’UTR-Mt*^) constructs. Changes of bases on 5’UTR sequence in the mutant construct (asterisk) are described in method sections. HEK293 cells were co-transfected with Flag-tagged STAU1, and wildtype or mutant *5’UTR-MTOR-LUC* or empty LUC reporter constructs followed by luciferase assay (**F, G**). No *s*ignificant changes in luciferases activity are achieved by mutant *MTOR*^*5’UTR-Mt*^ construct. Luciferase expression was normalized to Renilla luciferase (RLU). Results are from three replicate experiments. ACTB was used as a loading control, and representative blots of three independent experiments are shown. One-way ANOVA followed by Bonferroni’s multiple comparisons test. Data are mean ± SD, ns = *P* > 0.05, *\*\*\*P* < 0.001

### STAU1 overabundance impairs autophagic flux

Increases in LC3-II abundance by western blotting (**Fig 1**)^17^ are good indicators of changes in autophagy but alone do not determine increased on-rates due to expansion of the autophagosome pool vs. decreased off-rates due to reduced autophagic flux. We used three independent assays to examine autophagic flux. First, we employed bafilomycin A1 (Baf), an inhibitor of lysosome-autophagosome fusion. As expected, Baf treatment of wildtype cells increased GFP-LC3-II levels in a dose-dependent manner. In ATXN2-Q58 KI cells^11^, however, Baf treatment did not further increase GFP-LC3-II levels, consistent with blocked autophagosome-lysosome fusion **(Fig 5A, B)**. In a second cellular model using C9orf72 patient fibroblasts, LC3-II levels likewise did not increase with increasing Baf doses **(Fig 5C, D)**. These observations in both cell lines are consistent with decreased autophagic flux.

**Figure 5.**
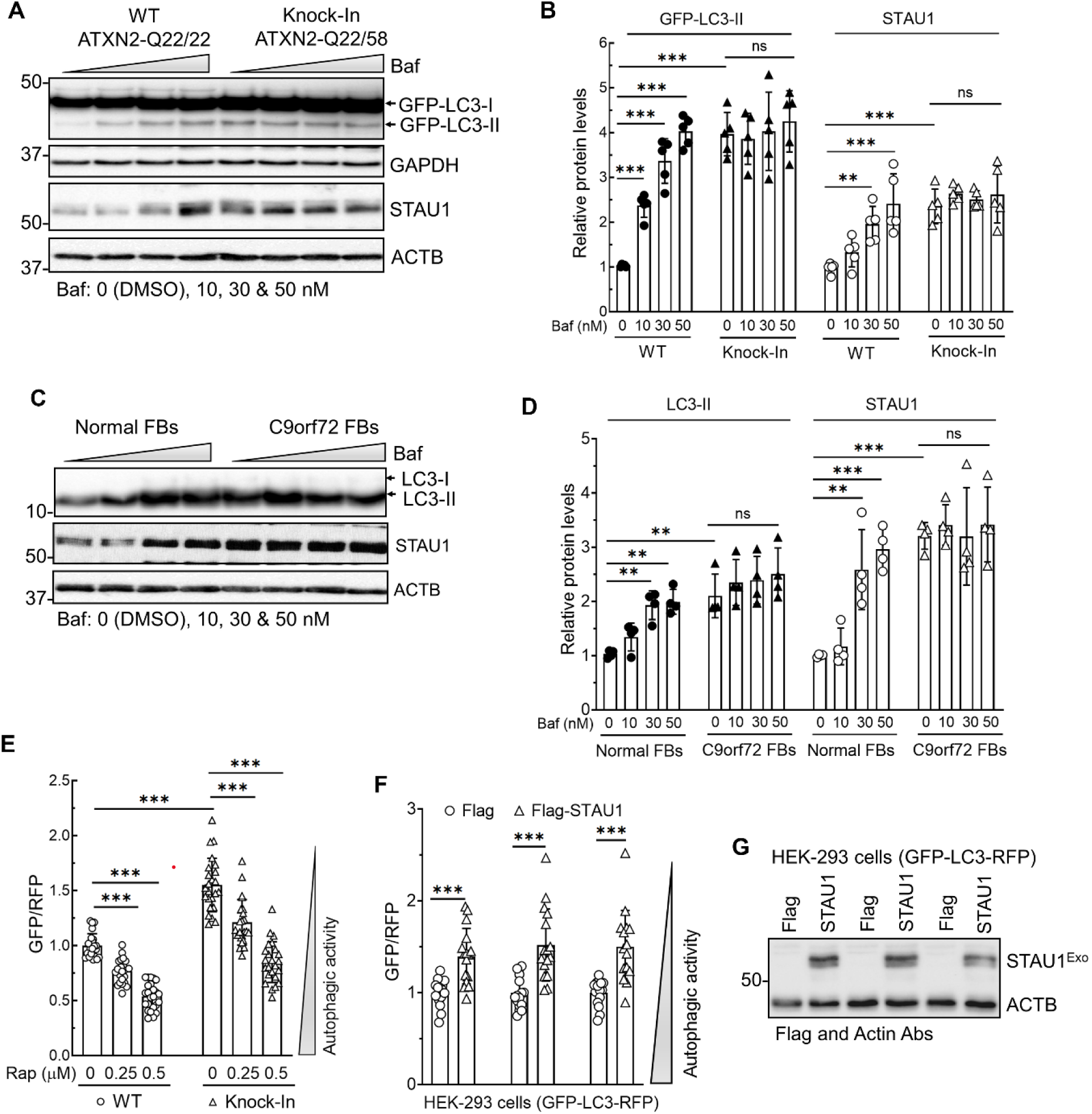
Autophagic flux is impaired in SCA2 and C9orf72 cells. **(A)** HEK293 and ATXN2-Q58 KI cells, stably expressing GFP-LC3, were treated dose-wise with Bafilomycin A1 (Baf) for 6 h and analyzed by western blotting. As expected, increasing doses of Baf increased GFP-LC3-II levels by blocking auto-lysosome fusion in wildtype HEK293 cells, whereas no further increase is seen in cells with mutant ATXN2, consistent with reduced autophagic flux. Baf treatment also shows increased STAU1 levels in wildtype cells without affecting in mutant cells. **(B)** Quantification of western blots in (**A**) (n=5 blots). All proteins were normalized to GAPDH or ACTB. **(C, D)** C9orf72 FBs show inefficient autophagosome-lysosome fusion. Normal and C9orf72 FBs were treated dose-wise with Baf for 6 h and analyzed by western blotting. Baf treatment showing increased levels of LC3-II and STAU1 levels in normal FBs without affecting those levels in C9orf72 FBs (**C**). **D)** Quantification of LC3-II and STAU1 levels on western blots. **(E)** Autophagic flux measured using the dual-color, cleavable GFP-LC3-RFP reporter. ATXN2-Q58 KI cells stably expressing the reporter showed a significantly increased GFP:RFP ratio compared to wildtype cells, indicating low autophagic flux. Treatment with rapamycin (Rap) for 24 h, an inhibitor of mTOR and autophagy inducer, improves autophagic flux in a dose-dependent fashion. **(F, G)** STAU1 expression lowers autophagic activity. HEK293 cells, stably expressing GFP-LC3-RFP, were transfected with Flag or Flag-tagged STAU1 (3 independent transfections) for 48 hr followed by measuring GFP:RFP ratio **(F)** and western blot analyses **(G)** showed a significantly increased GFP:RFP ratio. ACTB was used as loading control and representative blots of three independent experiments are shown. Ordinary one-way ANOVA followed by Bonferroni’s multiple comparisons test. Data are mean ± SD, ns = *P* > 0.05, *\*P* < 0.05, \*\**P* < 0.01, *\*\*\*P* < 0.001

In a 2^nd^ assay, we used an ATG4-cleavable dual GFP-LC3-RFP reporter^30^. Following cleavage by ATG4, GFP-LC3 undergoes lipidation and is degraded by autophagy, while RFP remains in the cytosol, serving as an internal control. Under conditions of decreased autophagic flux, autophagic GFP-LC3 degradation is decreased resulting in an increased GFP/RFP ratio^30^. As shown in **Fig 5E**, flux is significantly reduced in ATXN2-Q58 KI cells compared to wildtype HEK293 cells. Upon treatment with rapamycin, the abnormally elevated GFP/RFP ratio was reduced, consistent with normalization of autophagic flux **(Fig 5E)**. In the 3^rd^ experiment, we tested whether STAU1 overabundance in itself was sufficient to inhibit autophagic flux generating wildtype HEK293 cells stably expressing the GFP-LC3-RFP reporter. Upon exogenous expression of STAU1 in this cell line, flux was significantly impaired **(Fig 5F, G)**. These results indicate that STAU1 overabundance in itself is sufficient to reduce autophagic flux via mTOR signaling.

### Inhibition of mTOR activity lowers STAU1 levels

To further define the relationship between STAU1, *MTOR* mRNA, and autophagy, we sought to manipulate mTOR signaling using the mTOR inhibitor^3,5^, rapamycin, and siRNA to *MTOR*. When HEK293 or ATXN2-Q58 KI cells were treated with rapamycin for 24 hrs, a significant reduction of STAU1 abundance was seen in ATXN2-Q58 KI cells accompanied by reduced levels of p62 and LC3-II. However, rapamycin treatment in control HEK293 cells did not affect the levels of STAU1 and autophagic pathway protein components **(Fig 6A, B)**. Similar results were obtained using *MTOR* RNAi **(Fig 6C, D)**. Thus, reducing STAU1 levels via mTOR inhibition rescued the toxic effect of mutant ATXN2 on autophagy observed in SCA2.

**Figure 6.**
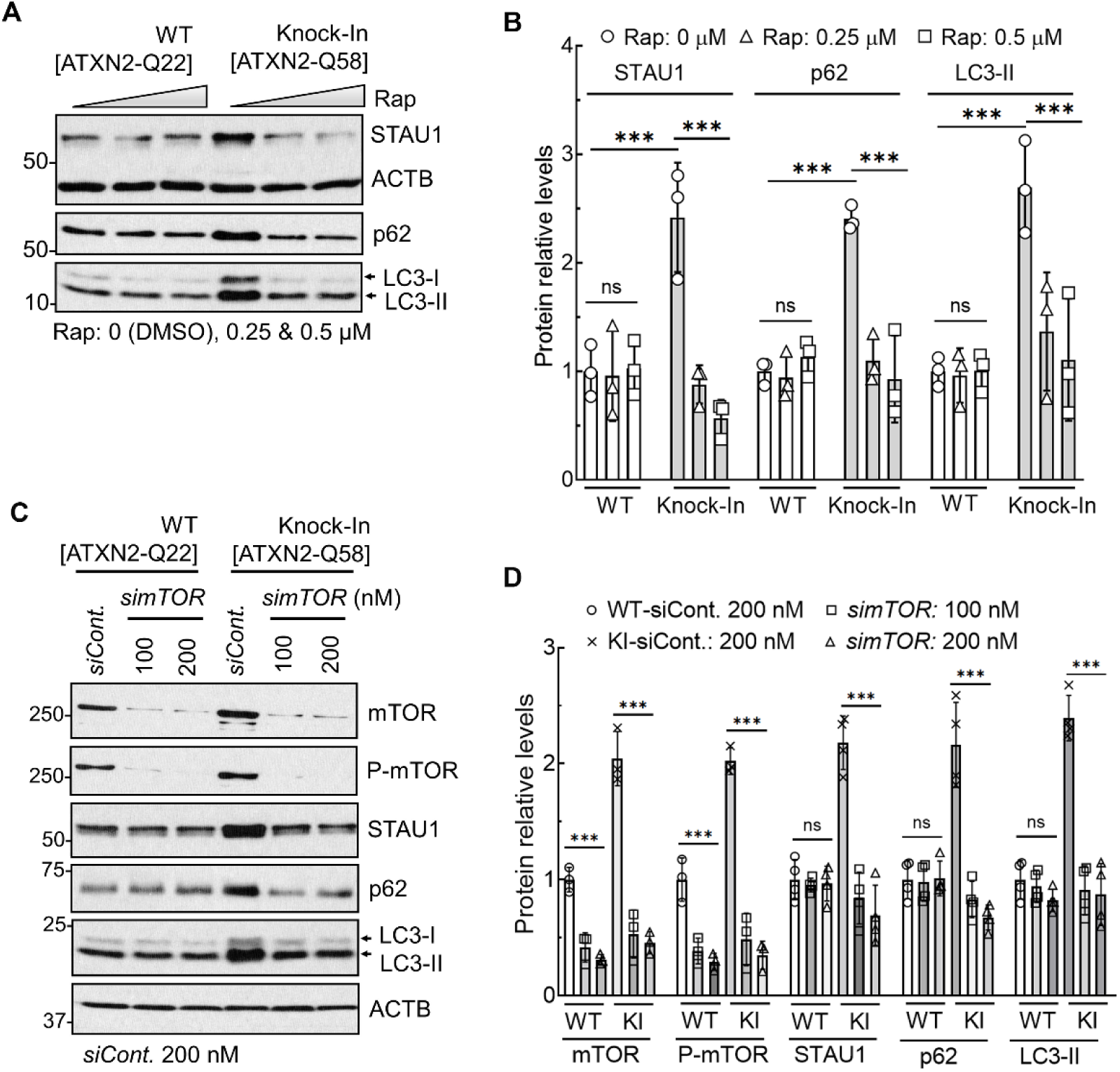
mTOR inhibition improves STAU1 clearance. **(A, B)** HEK293 and ATXN2-Q58 KI cells were treated dose-wise with Rapamycin (Rap) for 24 h followed by western blotting. Rap-induced mTOR inhibition lowers STAU1 levels along with p62 and LC3-II in ATXN2-Q58 KI cells without affecting those levels in wildtype cells. **(B)** Quantification of western blots in (**A**). **(C, D)** RNAi-mediated mTOR depletion lowers STAU1 and restores autophagy marker proteins. HEK293 and ATXN2-Q58 KI cells were transfected with *MTOR* RNAi, and cells were analyzed by western blotting at 4 days post-transfection. *MTOR* RNAi-mediated mTOR and phospho-mTOR depletion showing reduced STAU1 and autophagic pathway protein component levels in ATXN2-Q58 KI cells without affecting those levels in wildtype cells. **(D)** Quantification of western blots in (**C**). Ordinary one-way ANOVA followed by Bonferroni’s multiple comparisons test. Data are mean ± SD, ns = *P* > 0.05, *\*\*\*P* < 0.001. ACTB was used as loading control and representative blots of three independent experiments are shown

### STAU1 depleted cells are resistant to poly-GA aggregates and mTOR activation

ALS and FTD with the intronic *C9orf72* G_4_C_2_ expansion results in the expression of dipeptide repeat (DPR)/repeat-associated non-AUG (RAN) proteins [poly-(GA, GP, GR, PR, and PA)] and shows predominantly neuronal cytoplasmic inclusions in C9orf72 patients^31-33^. As STAU1 overabundance and mTOR hyperactivity are evident in C9orf72 FBs and an animal model (**Fig 1**), we tested whether DPRs by themselves would change STAU1 and mTOR levels. Of the 5 DPRs generated by RAN-translation, we chose to analyze GA- or PA-DPRs owing to their significant toxicity^33-36^. The exogenous expression of Flag-tagged GA^50^ or PA^50^ in HEK293 cells caused increased STAU1 levels as well as mTOR and its downstream targets (**Fig 7A**). We also observed poly-GA and poly-PA protein aggregations (high molecular weight species), however, poly-GA protein showed a tendency for aggregation within the stacking gel on western blot (**Fig 7A**), which was consistent with previous experimental observations in cultured cells^37,38^.

**Figure 7.**
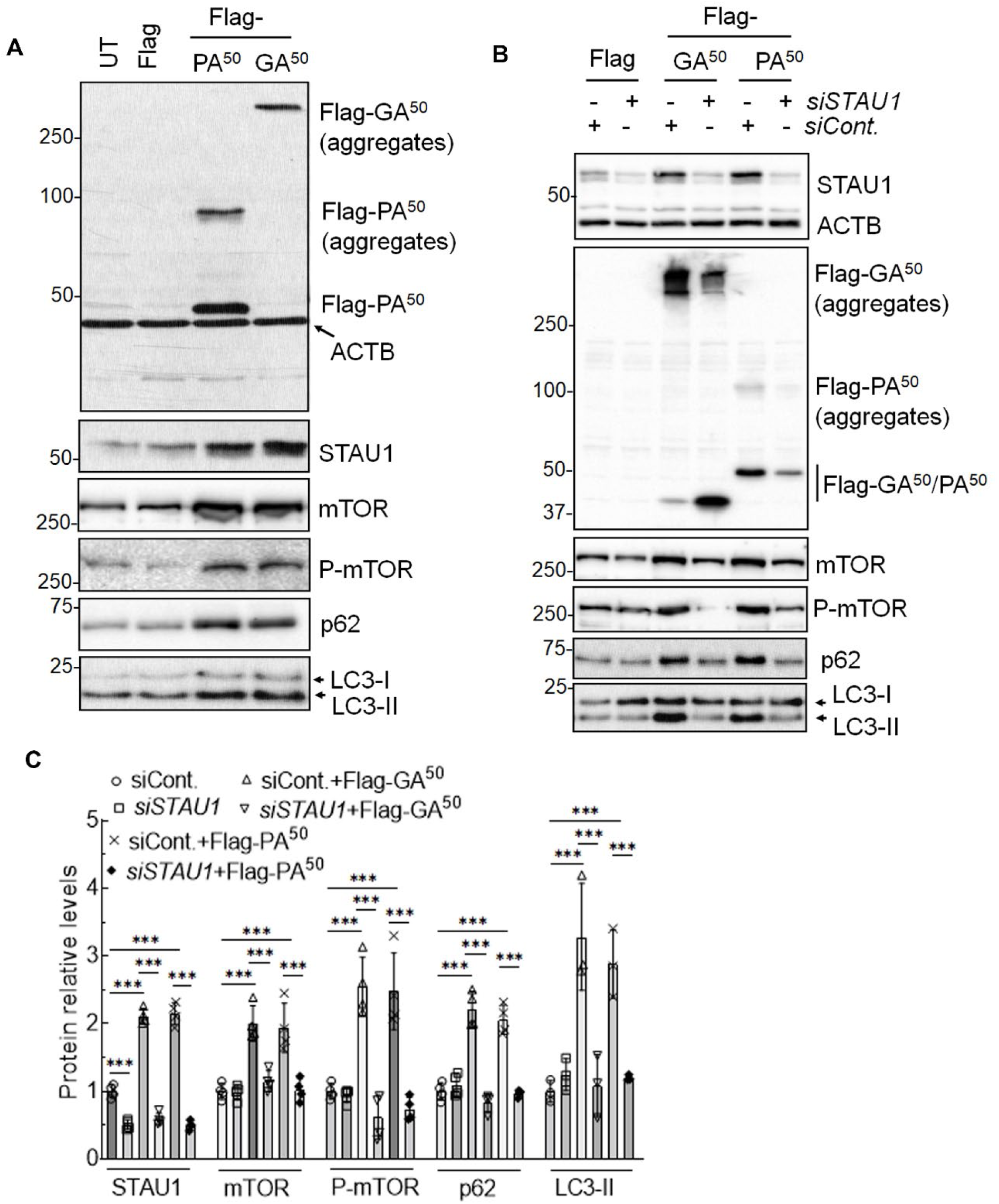
STAU1 depleted cells are resistant to DPR aggregates and mTOR activation. **(A)** DPR expression in cultured cells associated with STAU1 overabundance and mTOR activation. HEK293 cells were transfected with Flag-tagged GA^50^ or PA^50^ plasmid constructs for 48 hrs followed by western blotting. Both Poly-GA and poly-PA showed aggregation (high molecular weight species), with poly-GA protein detected as aggregates form within the stacking gel on western blot. Expression of exogenous Flag-tagged poly-GA or poly-PA resulted in increased STAU1 levels and mTOR activation. **(B, C)** HEK293 cells were treated 72 hrs by *STAU1* RNAi and then transfected with Flag-tagged GA^50^ and PA^50^ plasmids for an additional 48 hrs followed by western blotting. STAU1-depleted cells have reduced poly-GA aggregates and poly-PA protein in abundance, and resistance to mTOR activation upon overexpression of DPRs compared with controls. ACTB was used as loading control, and representative blots of 3 independent experiments are shown. **(C)** Quantification of western blots in (**B**). Ordinary one-way ANOVA followed by Bonferroni’s multiple comparisons test. Data are mean ± SD, ns = *P* > 0.05, *\*\*\*P* < 0.001. ACTB was used as loading control, and representative blots of 3 independent experiments are shown.

Since lowering STAU1 levels can normalize aberrant autophagy pathway proteins associated with SCA2, TDP-43, and C9orf72 mutations^17^, we wanted to test if reducing STAU1 levels were protective of DPR-induced pathological stress. We treated HEK293 cells with *STAU1* RNAi followed by transfection of plasmids encoding DPRs [poly-GA or poly-PA] (**Fig 7B**). Lowering STAU1 levels reduced aggregated poly-GA, increased non-aggregated poly-GA, and decreased aggregated with non-aggregated forms of poly-PA (**Fig 7B, C**). This also resulted in normalization of mTOR signaling, and p62 and LC3-II levels (**Fig 7B, C**). These results establish that STAU1 plays a key role in protecting cells from DPR-mediated pathological stress.

### Reduced Staufen1 levels ameliorate autophagy function across animal disease models

The increased abundance of STAU1 in NDDs and its association with dysregulation of autophagic flux led us to predict that STAU1 can be directly targeted to alter autophagy function. Previously, we demonstrated that *Stau1* haploinsufficiency partially restored behavioral, proteomic, and morphological characteristics of SCA2 pathology *in vivo*^11^. In accordance with this, in *ex vivo* STAU1 reduction via RNAi also showed normalization of mTOR pathway proteins in SCA2, ALS-TDP-43, and C9orf72-FTD cellular models^17^. These data indicate that reducing STAU1 levels may be useful for preventing maladaptive responses to cellular stress observed in NDDs.

### In vivo results

The availability of a mouse with a *Stau1* loss-of-function allele allowed us to determine the effect of Stau1 reduction using genetic interaction^21^. For these experiments, we used two well-established mouse models of neurodegeneration that model human diseases, the *Pcp2*-*ATXN2*^*Q127*^ SCA2 mouse, and the *Thy1*-TDP-43 ALS/FTD mouse^11,20,22^. We tested restoration of autophagic activity by measuring the levels of mTor, P-mTor, P-S6K, p62, and LC3-II by quantitative western blot analysis. Lowering *Stau1* in wildtype animals did not alter autophagy marker proteins in an appreciable way (**Fig 8A, B, C, D**). In contrast, reducing pathologically elevated Stau1 in SCA2 mice improved markers of autophagy in a dose-dependent fashion such that levels of all proteins were returned to near normal (**Fig 8A, B, C, D**). Next, we assayed if genetic reduction of *Stau1* in *TDP-43*^*Tg/+*^ mice (22 wks of age) reduce mTOR activity^21,22^. Similar to SCA2 mice, reducing *Stau1* genetic dosage by only 50% in *TDP-43*^*Tg/+*^ mouse spinal cord extracts resulted in decreased mTOR activity and restoration of autophagy pathway proteins (**Fig. 8E, F)**. Taken together, these data demonstrate that lowering Staufen1 levels restores autophagy function in SCA2 and ALS/FTD -TDP-43 animal models.

**Figure 8.**
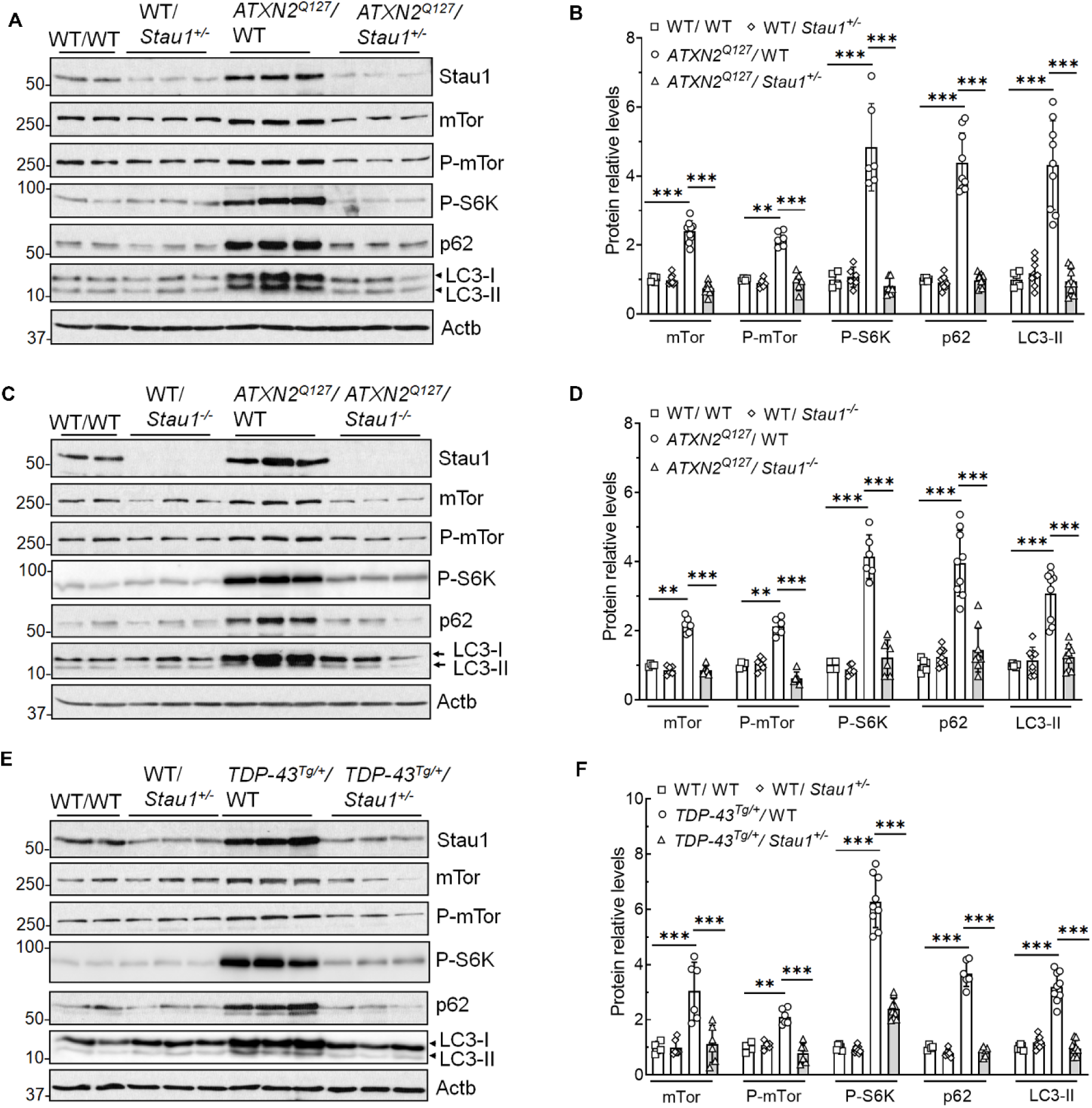
Reduction of Stau1 levels *in vivo* attenuates mTor activation in SCA2 and TDP-43 ALS models. **(A-F)** *ATXN2*^*Q127*^ mice with haploinsufficiency or absence of *Stau1* restores autophagic pathway proteins. Western blotting of cerebellar extracts from a cross of *ATXN2*^*Q127*^*/Stau1*^*+/-*^ mice (34 wks of age)^11^ (**A**) or a cross of *ATXN2*^*Q127*^*/Stau1*^*-/-*^ mice (20 wks of age) (**C**) showing reduced mTor activity and normalization of autophagic pathway proteins *in vivo*. **(E)** Western blotting of spinal cord extracts from *TDP-43*^*Tg/+*^ hemizygous mice (22 wks of age) haploinsufficient for *Stau1* showing reduced mTor activity and normalization of autophagic pathway proteins. Each lane represents extract from an individual mouse and 3 animals per group. Actb was used as a loading control and the blots are from three replicate blots. **(B, D, F)** Quantitative analysis of western blots in **(A, C, E)** demonstrating near normalization of mTOR pathway proteins. Ordinary one-way ANOVA followed by Bonferroni’s multiple comparisons test. Data are mean ± SD, ns = *P* > 0.05, \*\**P* < 0.01, *\*\*\*P* < 0.001

## Discussion

Identifying protein interactors has been an important approach to understanding pathways altered by the presence of mutant disease proteins. A connection to ALS was found by showing that ATXN2 interacted with the RNA-binding protein TDP-43 which has been implicated in multiple NDDs^39^. This prompted us to determine other ATXN2 interacting proteins. Using immunoprecipitation coupled with mass spectrometry, we found that STAU1 precipitated with ATXN2 with no differences in strength of interaction between wildtype and mutant ATXN2^11^.

Staufen is a well-studied protein in fly development^40^. Through gene duplication, two Staufen genes exist in mammalian cells, whereby Staufen2 (STAU2) is more similar to fly Staufen than STAU1. STAU1 and STAU2 are a dsRBP. They are found in stress granules (SGs) in brain oligodendrocytes and neurons and modulate SG dynamics^13,41-43^. Owing to Staufen’s ability to function as an RBP, it is involved in regulating RNA metabolism, mRNA transport in neuronal dendrites, and other cells in vertebrates^44-46^. Furthermore, STAU1 regulates the translational efficiency via 5’UTR and polysome association, and the stability of specific transcripts through their 3’UTRs, a mechanism referred to as STAU1-mediated RNA decay (SMD)^15,16,26,47^.

Mutations in heterogeneous nuclear ribonucleoproteins and TIA1, a SG protein, can cause ALS and FTLD^48,49^. Direct mutations in *STAU1* have not been described, but our observations of STAU1 overabundance in several neurodegenerative disease models, patients with *C9orf72* expansion, and sporadic ALS support the hypothesis of STAU1’s role in neurodegeneration^17^. It is possible that STAU1 is more directly involved in autophagic feedback regulation in aggregate-prone disease proteins. Although we have shown that STAU1 is increased in a number of NDDs, recent comparisons of SCA1 and SCA2 mouse models showed different states of autophagy^50^. Additional work will be required to characterize the role of STAU1 in abnormal autophagy and the potential function of STAU1 in clearance of toxic protein aggregates. It is tempting to speculate that *STAU1* variants will be identified in ALS or FTD patients as mendelian or risk alleles.

Our results have implications for the basic understanding of mechanisms regulating autophagy and for identifying novel targets for the treatment of neurodegeneration. Current understanding of autophagy emphasizes regulation by kinase cascades and by transcriptional changes in key genes involved in autophagy^1-2,51^. A number of RBPs other than autophagy signaling pathway components like mTOR are involved in regulating autophagy-related mRNA metabolism such as maintaining their stability, modulating their translation, and regulating splicing^52^. We now describe an additional regulatory mechanism that involves RNA-binding proteins like STAU1 with the ability to directly influence the translation of already transcribed mRNAs. Although this process may be homeostatic in the setting of acute stress, it is likely maladaptive with chronic neuronal stress. We showed that STAU1 overexpression in HEK293 cells increased mTOR levels and activity and that STAU1 directly interacted with *MTOR* mRNA and enhanced its stability leading to increased translation. Consistent with reduced autophagic flux, STAU1 overexpression caused increased p62 and LC3-II abundance (**Fig 3**)^17^.

As STAU1 is mainly degraded by autophagy, direct interaction of STAU1 with the 5’-UTR of mRNAs encoding mTOR could result in a deleterious feed-forward loop in that elevated mTOR further decreases autophagic flux. As we have shown previously, elevated STAU1 also affects SMD with the potential to alter neuronal transcriptomes significantly as well as the UPR and potentially DPR clearance through multiple mechanisms. Finally, our results suggest that multiple different stressors can result in STAU1 elevation with the potential that this maladaptive response can be engaged by genetic, infectious, or environmental stressors **(Fig 2)**.

### STAU1 in translation regulation and the role of RNA binding domain

The double-stranded RNA-binding domains (dsRBDs) of STAU1 and dsRNA-stem loop interaction are critical to mediate STAU1 interactions with target mRNAs. The RBDs are highly conserved in Staufen homologues from *Drosophila* to humans. Of the four dsRBDs in STAU1, the dsRBD3 acts as the major functional domain for target RNA interactions^28,29^. RNAs containing >8 bp of dsRNA bind to dsRBD3, but optimal binding is observed with RNAs of 12 bp or longer double-helical stems^28^. Mutations in the dsRBD3 or disruption of the helical structure of target RNAs by unpaired bases significantly abolish STAU1 and RNA interactions^28^. These findings parallel the observation of dsRBDs of RNA-activated protein kinase R (PKR)^53,54^. STAU1 mainly recognizes flexible secondary structures rather than specific consensus sequences. It is accompanied by a multistep process: STAU1 dsRBD3 recognizes and scans dsRNA stem-loops, allowing the recruitment of dsRBD4 on the Staufen binding site (SBS). dsRBD4 then finalizes contacts with the RNA and allows dsRBD3 to position correctly^55^. Consistent with this, we observed that mTOR levels remained unchanged when the dsRBD3 was deleted (**Fig 3**). On the other hand, when the *MTOR* 5’UTR RNA stem-loop structure was disrupted by changing non-pairing bases, STAU1 showed reduced interaction towards *MTOR* - 5’UTR (**Fig 4**). It is likely that other RBDs and even RNAs will influence specific RNA binding *in vivo*.

### Challenges of autophagic flux evaluation

Measuring p62 is useful to evaluate autophagic flux. p62 binds to LC3 and incorporates into autophagosomes serving as a substrate of autophagy^56,57^. The total cellular p62 expression levels inversely correlate with autophagic activity in that p62 stabilization reflects autophagy suppression. Of note, p62 positive aggregates are common in NDDs including C9orf72 associated disease^58^. Consistent with this, p62 stabilization across neurodegenerative cellular and mouse models and upon STAU1 overexpression (**Figs 1 and 3**)^17^ support the mechanistic role for STAU1 in modulating autophagic flux in neurodegeneration.

Previously, we demonstrated that STAU1 was overabundant in neurodegenerative disease FBs, mouse models, and human tissues and that overabundance was associated with mTOR hyperactivation^17^. We now show Stau1 and mTor abundances in the ALS/FTD BAC-*C9orf72* mouse **(Fig 1)**. Associated with elevated STAU1 and mTOR, we also found increased levels of p62 and LC3-II predicting reduced flux by inefficient autophagosome-lysosome fusion (**Fig 1**). Increased autophagosome formation is known to be associated with increased levels of a phosphatidylethanolamine (PE)-conjugated form of LC3-II^59^. However, not all LC3-II is present on autophagic membranes, and a subpopulation of LC3-II can be ectopically generated in an autophagy-independent manner^60^. Therefore, care must be taken in interpreting increases in LC3 levels as this may occur as a result of an increase in autophagosome formation (autophagic activity) or autophagosome stabilization due to defects at downstream steps (e.g., autophagosome-lysosome fusion)^56,57^. To differentiate between these conditions, we used a GFP-LC3 fluorescence probe to assess autophagic flux. As expected, treatment with bafilomycin A1 (Baf), a lysosomal inhibitor, increased GFP-LC3-II and STAU1 levels in wildtype cells in a dose-dependent manner. In ATXN2-Q58 KI cells, however, Baf treatment did not further increase GFP-LC3-II and STAU1 levels compared to untreated controls, reflecting reduction of autophagosome-lysosome fusion (indicating reduced autophagic flux) (**Fig 5**)^56-57^. These results are also consistent with our observations in C9orf72 fibroblasts upon Baf treatment (**Fig 5**).

The potential caveat is that GFP-LC3 lacks an internal control. Given this potential limitation, we used a dual-fluorescence probe, GFP-LC3-RFP, to reassess autophagic flux^30^. When expressed in cells, the GFP-LC3-RFP protein is cleaved by endogenous ATG4 to yield equimolar amounts of GFP-LC3 and RFP. GFP-LC3 undergoes lipidation and attaches to the autophagosome membrane and is quenched/degraded by autophagy. Meanwhile, RFP stays in the cytosol and serves as an internal control. The GFP/RFP signal ratio reversely correlates with autophagic activity, *i*.*e*., an increasing GFP/RFP ratio represents reduced autophagy flux^30^. As the expression of this probe does not affect autophagic flux and mTORC1 activity^30^, this probe is well suited as a flux assay. Using the GFP-LC3-RFP probe, ATXN2-Q58 KI cells had higher GFP/RFP ratios than wildtype HEK293 cells, reflecting reduced autophagic flux (**Fig 5**). Reduced flux was reversible, as treatments with the mTOR inhibitor rapamycin resulted in a decreased GFP/RFP ratio in a dose-dependent manner. Using the GFP-LC3-RFP probe, we also showed that STAU1 overexpression by itself in wildtype HEK293 cells was sufficient to reduce autophagic flux (**Fig 5**).

### Restoration of STAU1 by rapamycin or MTOR RNAi

Our study shows association of mTOR activation and reduced autophagic flux with STAU1 overabundance. HEK293-ATXN2-Q58 KI cells treated with rapamycin had increased autophagic flux **(Fig 5)**. Autophagy induction by rapamycin has resulted in clearance of aggregate-prone disease proteins, including mutant huntingtin protein^61^, and TDP-43 in FTD^3^. This parallels our observation for restoration of STAU1 and autophagy pathway protein levels in our SCA2 cellular model upon rapamycin treatment or lowering mTOR expression by *MTOR* RNAi **(Fig 6)**. Although rapamycin promotes autophagosome formation^62^, induction of autophagosome formation without parallel enhancement of autophagic flux may result in further accumulation of autophagosomes and more neuronal damage. Targeting STAU1 however appears to both restore autophagosome abundance and normalize autophagic flux.

### DPR role in cell death and autophagy

Transcripts with expanded G_4_C_2_ tracts in *C9orf72* are RAN-translated into toxic DPR proteins that are derived by different reading frames from both strands: poly-GA, poly-GR, poly-GP, poly-PR, and poly-PA^31-33^. These have been detected as neuronal cytoplasmic aggregates in patient brain frontal cortex or cerebellum^32-36,63^. Although arginine-containing DPRs (poly-GR and poly-PR) are more toxic, poly-GA is the most abundant of the DPRs in patients^35,63^. Similar to poly-GA toxicity in cell culture models, mice overexpressing poly-GA exhibit motor and cognitive deficits combined with cerebellar atrophy, astrogliosis, TDP-43 pathology, and neuronal loss^64-66^, demonstrating the important role of poly-GA’s in C9orf72 ALS-FTD pathogenesis. Thus, formation of DPR aggregates could be attributed to the autophagy malfunction in C9orf72-FTD as aggregates are degraded by autophagy^63^. Our data show the association of STAU1 and mTOR abundances upon poly-GA and poly-PA expression and aggregation (higher molecular weight species) of poly-GA and poly-PA proteins on immunoblots at the molecular level (**Fig 7**). This very high molecular weight poly-GA aggregate nature is consistent with other published observations^37,38^. One possible explanation for this may be due to hydrophobic nature of poly-GA and its tendency to self-aggregate and run as high molecular weight species on blots. An additional putative mechanism for STAU1-modulation of DPR abundance and aggregation **(Fig 7)** is through its bidirectional control of eIF2α phosphorylation status, which regulates cap-dependent translation and the activation of the integrated stress response (ISR). We have previously shown that STAU1 overabundance increased phospho-eIF2α, whereas STAU1 depletion led to a decrease, associated with attenuation of ER stress and apoptosis in cellular models, including SCA2, ALS-TDP-43, and FTD-C9orf72^18^. This favors association of STAU1, ISR activation, and DPR expression in FTD-C9orf72. Because activation of the ISR triggered by phospho-eIF2α drives RAN translation of DPRs, while its suppression reduces DPR levels in multiple models^67,68^, we hypothesize STAU1 could regulate DPR abundance via its actions on phospho-eIF2α. In line with this, we show that poly-GA and poly-PA are less aggregated when STAU1 is depleted from cells (**Fig 7**). Thus, targeting STAU1 could counteract the accumulation of DPRs in C9orf72-FTD.

### Reduced Staufen1 levels ameliorate autophagy function in vivo

Due to the fundamental role of autophagy in maintaining homeostasis, its inhibition has deleterious effects in both healthy and neurodegenerative states. Our findings that STAU1 overabundance causes mTOR activation in neurodegenerative cell and animal models are consistent with other observations^52^. To prevent adverse effects of STAU1, it is key to determine whether *STAU1* silencing improves autophagy flux. Reducing STAU1 levels by RNAi resulted in normalization of mTOR activity in SCA2, ALS-TDP-43, and FTD-C9orf72 cellular models^17^. Our *in vivo* genetic interaction experiments in SCA2 mice showed that *Stau1* haploinsufficiency reduced ATXN2 aggregates in PCs and improved motor function^11^. Like mutant ATXN2 aggregates in SCA2, TDP-43 aggregates are a common neuropathological feature of ALS and FTD, including C9orf72 disease^63^. TDP-43 also aggregates with p62 resulting in p62 stabilization, which is an indicator for defective autophagy^58,63^. Consistent with this, genetic reduction of *Stau1* in ATXN2^Q127^ SCA2 mice showed improved markers of autophagy in a dose-dependent fashion such that levels of all proteins were returned to near normal. Cerebellar and spinal cord tissues from transgenic *TDP-43*^*Tg/+*^ mice have impaired autophagy. Reducing *Stau1* by only 50% through genetic interaction resulted in decreased mTOR activity and restoration of autophagy pathway proteins (**Fig 8**). Further studies will be needed to replicate these findings in other TDP-43 mouse models including analysis of improved neuronal survival and behavior.

In summary, we demonstrated that STAU1 overabundance drives mTOR hyperactivation in the presence of disease mutations. STAU1 may be an attractive novel target for antisense oligonucleotide (ASO)- or miRNA viral-based therapies for neurodegenerative phenotypes *in vitro* and *in vivo*. For C9orf72/FTD-DPR associated phenotypes, targeting STAU1 may provide an additional target as the *C9orf72* gene product appears to have essential cellular functions.

## Acknowledgments

This work was supported by National Institutes of Neurological Disorders and Stroke (NINDS) grants R37NS033123 and R56NS33123 to SMP, R01NS097903 to DRS, and R21NS103009 and R21NS081182 to DRS and SMP. DRS was also supported by a Harrington Discovery Institute Rare Disease Scholar award. The funders had no role in study design, data collection and analysis, decision to publish, or preparation of the manuscript.

## Author contributions

S.P., W.D., K.P.F., D.R.S., and S.M.P. conceived and designed the experiments. Experiments were performed by S.P., W.D., M.G. and K.P.F. BAC-*C9orf72* mouse brain extracts are provided by T.Z. and L.P.W.R. Data analyses were performed by S.P., W.D., M.G., K.P.F and D.R.S. and S.M.P. All figures were generated by S.P. and W.D. Administrative assistance was provided by K.P.F. The manuscript was written by S.P., W.D., K.P.F., M.G., D.R.S., and S.M.P., and critically reviewed by L.P.W.R., D.R.S., and S.M.P.

## Potential conflicts of interest

The authors declare no competing interests.

## Data availability

The data that support the findings of this study are included in this article.

## Notes

### Competing Interest Statement

The authors have declared no competing interest.

